# FcyRIIB is a novel immune checkpoint in the tumor microenvironment limiting activity of Treg-targeting antibodies

**DOI:** 10.1101/2023.01.19.522856

**Authors:** David Knorr, Rom Leidner, Shawn Jensen, Ryan Meng, Andrew Jones, Carmen Ballesteros-Merino, R. Bryan Bell, Maria Baez, David Sprott, Carlo Bifulco, Brian Piening, Rony Dahan, Bernard A. Fox, Jeffrey Ravetch

**Affiliations:** Laboratory of Molecular Genetics and Immunology, Rockefeller University, New York, NY; Department of Medicine, Memorial Sloan Kettering Cancer Center, New York, NY; Earle A. Chiles Research Institute, a division of Providence Cancer Institute, Portland, OR; Department of Immunology, Weizmann Institute of Science, Rehovot, Israel

## Abstract

Despite pre-clinical murine data supporting T regulatory (Treg) cell depletion as a major mechanism by which anti-CTLA-4 antibodies function in vivo, the two main antibodies tested in patients (ipilimumab and tremelimumab) have failed to demonstrate similar effects. We report analogous findings in an immunocompetent murine model humanized for CTLA-4 and Fcy receptors (hCTLA-4/hFcyR mice), where both ipilimumab and tremelimumab fail to show appreciable Treg depletion. Immune profiling of the tumor microenvironment (TME) in both mice and human samples revealed upregulation of the inhibitory Fcy receptor, FcyRIIB, which limits the ability of the antibody Fc fragment of human anti-CTLA-4 antibodies to induce effective antibody dependent cellular cytotoxicty/phagocytosis (ADCC/ADCP). Blocking FcyRIIB in humanized mice rescues Treg depleting capacity and anti-tumor activity of ipilimumab. For another target, CC motif chemokine receptor 8 (CCR8), which is selectively expressed on tumor infiltrating Tregs, we show that Fc engineering to enhance binding to activating Fc receptors, while limiting binding to the inhibitory Fc receptor, leads to consistent Treg depletion and single-agent activity across multiple tumor models, including B16, MC38 and MB49. These data reveal the importance of reducing engagement to the inhibitory Fc receptor to optimize Treg depletion by TME targeting antibodies. Our results define the inhibitory FcyRIIB receptor as a novel immune checkpoint limiting antibody-mediated Treg depletion in tumors, and demonstrate Fc variant engineering as a means to overcome this limitation and augment efficacy for a repertoire of antibodies currently in use or under clinical evaluation in oncology.

**Highlights:**

- Fully human anti-CTLA-4 antibodies are limited in their capacity to deplete T regulatory cells and drive durable anti-tumor immunity in humanized FcyR/hCTLA-4 mice
- The inhibitory Fcy receptor, FcyRIIB, is upregulated in the tumor microenvironment in patients and in humanized FcyR/hCTLA-4 mice
- Blocking FcyRIIB leads to rescue of Treg depletion in humanized murine models
- Fc engineering can improve the depleting capacity and in vivo anti-tumor activity of anti-CTLA and anti-CCR8 antibodies targeting tumor infiltrating Tregs

## Introduction

While cancer immunotherapy has revolutionized the treatment of various malignancies, factors leading to response and resistance remain poorly defined. Several cancer intrinsic markers such as high PD-L1 expression or tumor mutational burden have been used to predict response to both anti-PD-1 and anti-CTLA-4 immunotherapy.^1^ For CTLA4, several potential mechanisms of response are generally accepted. The most widely viewed mechanism of action is that antibody blockade of interactions of CTLA-4 with members of the B7 family removes this inhibitory checkpoint from effector T cells and promotes anti-tumor immunity.^2^ Recently, Waight and colleagues have proposed an Fc-dependent, but Treg depletion independent, mechanism of anti-CTLA-4 antibodies in which activating FcyRIIIA engagement leads to activation of antigen-specific T cells. While intriguing, these are primarily based on artificial in vivo stimulation assays (supertoxin co-administration) or in vitro studies using human PBMCs.^3^ Finally, based on several murine studies, depletion of intratumoral T regulatory cells expressing high levels of CTLA-4 in the TME removing immune suppression and allowing infiltration and expansion of tumor-specific CD8 T cells has been put forward.^4–6^ However, evidence for routine depletion of Tregs in patients receiving anti-CTLA-4 remains lacking, with recent data suggesting that both Ipilimumab and Tremelimumab fail to appreciably deplete Tregs in patients.^7^

Much of the research effort to determine whether human anti-CTLA-4 antibodies indeed deplete Tregs in vivo has relied on xenograft models, for which several caveats, including non-specific activation of immune cells, as well as differential expression of mouse and human FcyRs, are confounding factors.^8^ Arce Vargas and colleagues used humanized FcyR mice and chimeric anti-CTLA-4 antibodies (mouse Fabs with human Fcs) to determine contributions of the Fc fragment to in vivo activity.^5^ While these studies described the ability of human IgG1 antibodies to deplete Tregs in humanized models, they heavily relied on a tumor model (MCA-205) with robust immune infiltrate (ie >1-log fold higher infiltrating immune cells compared to MC38 and B16), not representative of most human tumor microenvironments. Additionally, while these studies also found that a combination of FcyRIIIA genotype (CD16a-V158F high affinity allele) and tumor mutational burden predicted favorable outcomes for patients treated with anti-CTLA-4 therapy, these variables fail to account for most patients responding to anti-CTLA-4 therapy.

Because of the disconnect between murine and human models of CTLA-4-mediated Treg depletion, and clear differences in the activating/inhibitory ratios between murine and human antibodies,^9^ we investigated whether FcyRIIB upregulation could potentially explain poor Treg-depleting capacity of Ipilimumab. Prior work has defined the inhibitory Fcy receptor, FcyRIIB, as an important regulator of ADCC/ADCP.^9,10^ FcyRIIB expression is known to be induced by several pathways, including IL-4 and HIF-1, both of which are readily found in the TME.^11,12^ Blockade of FcyRIIB has also been successfully employed to enhance ADCP of anti-CD20 antibodies,^13^ a process in which FcyRIIB competes with activating receptor engagement to limit ADCP, independent of ITIM-dependent FcyRIIB actvitity.^14^

We developed a fully humanized murine model to study Treg targeting antibody therapeutics with comparative analysis from human tissue specimens to determine the generality of the preclinical studies. Using this model system, we demonstrate increased FcyRIIB expression in the TME as a potential mechanism limiting therapeutic Treg depletion in patients by ipilimumab, tremelimumab, or newer second generation antibody approaches. Utilizing Fc engineering to decrease binding to the inhibitory FcyRIIB receptor and salvage anti-tumor efficacy, we demonstrate a novel strategy for Treg-targeted antibody therapeutics.

## Results

### Human anti-CTLA-4 antibodies fail to significantly deplete Tregs across tumor types in humanized CTLA-4/hFcyR mice

Recombinant human IgG antibodies have been a cornerstone of cancer treatment for almost two decades, and numerous studies have focused on clarifying the mechanisms by which anti-tumor immunotherapies elicit these therapeutic effects. As a result, the importance of Fcy receptor (FcyR)-mediated effector pathways for the elimination of tumors has been elucidated, resulting in the optimization of these interactions in second-generation anti-tumor immunotherapeutics.^15^ Antibodies have two ends, on one end the two identical Fab domains which recognize the target (e.g. CTLA-4, EGFR, HER2) while the other contains the Fc domain engaging myeloid effector cells. In order to determine the potential mechanisms by which anti-CTLA-4 antibodies deplete Tregs in vivo, we developed a murine line humanized for both human FcyRs and human CTLA-4 (hCTLA-4/hFcyR mice), allowing for evaluation of fully human antibodies in an immunocompetent setting. In brief, we used human CTLA-4 mice on a mouse CTLA-4-deficient background^6^ crossed to our previously described hFcyR mice (**Figure 1A**).^16^ The mice breed and develop normally without clinical evidence of any autoimmune phenomenon. The hCTLA-4/hFcyR mice express both hCTLA-4 and all human FcyRs in place of their murine homologues. When compared to WT C57BL/6 mice, hCTLA-4/hFcyR mice express similar levels of hCTLA-4 in both the tumor and secondary lymphoid tissues (**Figure 1B**). We next cloned the variable heavy and light sequences (Fabs) of iplimumab (clone 10D1) and tremelimumab (clone 1121), expressing each on IgG1 or IgG2 backbones for direct comparison of the contribution of both the Fab and Fc to their in vivo activity (**Figure 1C and Supplementary Figure 1A)**. Binding to human CTLA-4 was similar between the variants (**Figure 1D**), as was their ability to block interactions between human B7.1 and human CTLA4 (**Figure 1E**). When tested in the MC38 subcutaneous tumor model, which is responsive to murine or rat subclass anti-CTLA-4 antibody therapy, we found that 10D1-IgG1 led to a decreased proportion of Tregs compared to controls or 10D1-IgG2, and a reciprocal increase in infiltrating CD8 effector cells, consistent with prior reports (**Figure 1F**).^5,17^ To our surprise, ipilimumab and tremelimumab had similar in vivo efficacy in the MC38 colon carcinoma model (**Figure 1G**), including chimeric ipilimumab Fabs on the tremelimumab IgG2 Fc backbone (**Figure 1H**), suggesting no discernable differences in Fc contribution to activity in the MC38 model. We also tested different anti-CD4 subclass variants in the MC38 tumor model to assess whether lack of depletion is due to target-specific, rather than intratumoral effects. As expected, a decrease in the circulating CD4 T cell compartment was observed due to formation of immune complexes in the blood, but we did not see a decrease of intratumoral CD4 T cells, confirming that lack of cell-type specific depletion is not target-dependent, but a feature of the TME (**Supplementary Figure 1B**). We next assessed the ability of each ipilimumab human IgG subclass to deplete intratumoral Tregs in a more aggressive tumor model (B16 melanoma), where murine anti-CTLA-4 antibodies of the IgG2a subclass lead to Treg depletion and enhanced tumor control.^4^ Here, neither the 10D1-IgG1 or 10D1-IgG2 lead to changes in intratumoral CD8, CD4, or Tregs (**Figure 1I**), in line with what has been reported in patient samples. While modestly enhanced tumor control with 10D1-IgG1 treatment was initially observed, it did not lead to durable tumor control in the B16 model (**Figure 1J**).

**Figure 1.**
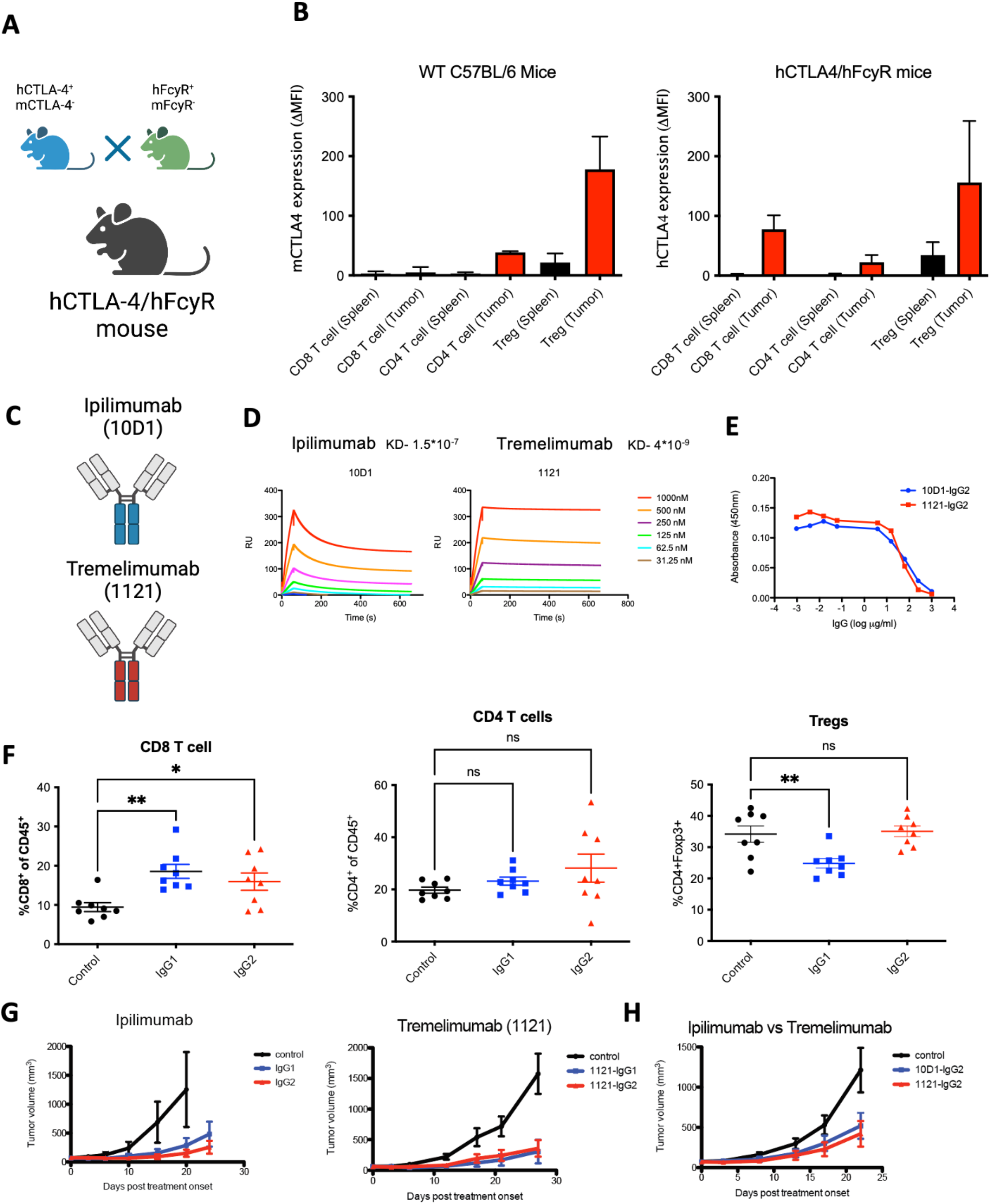
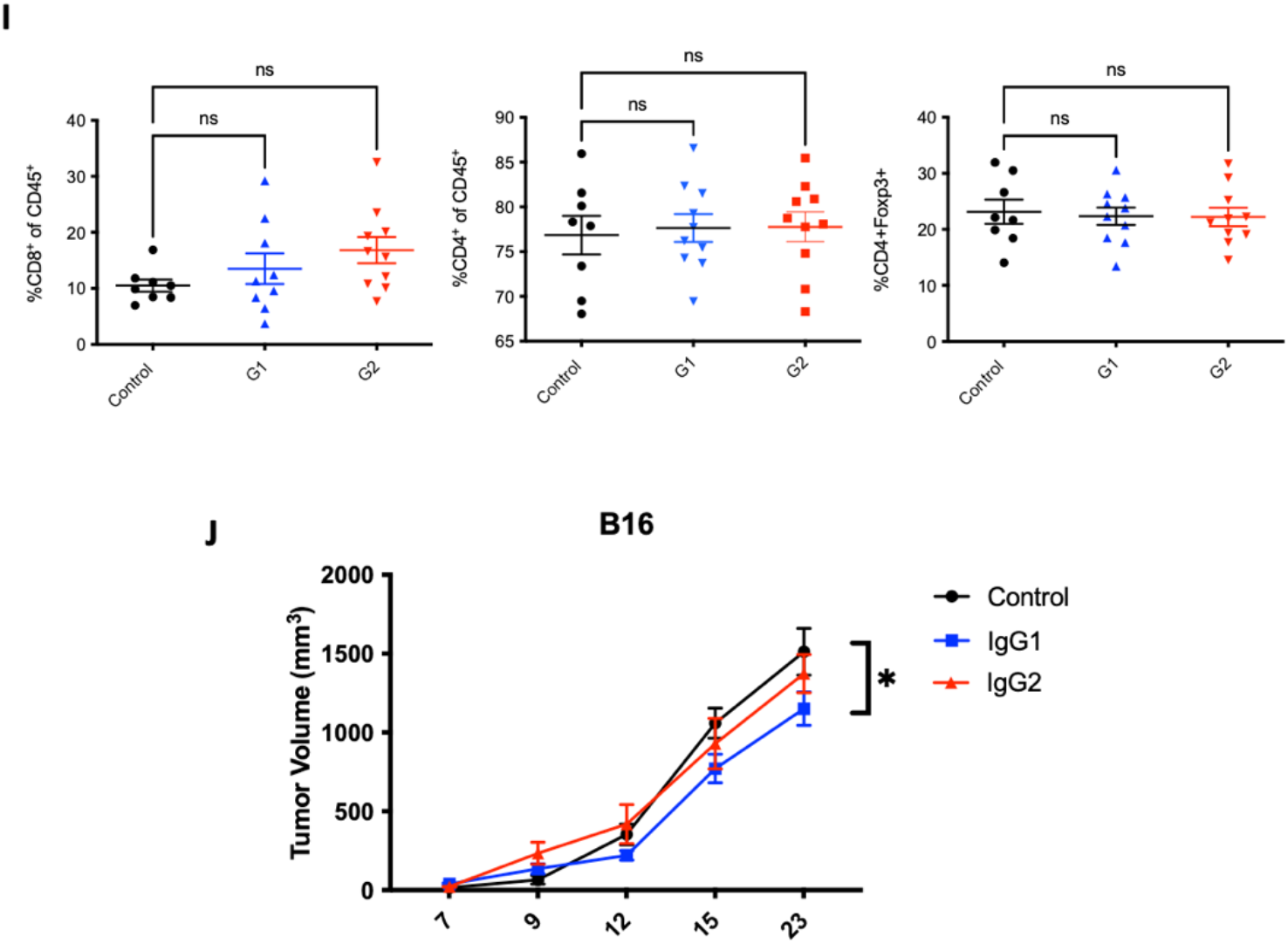
Development and characterization of ipilimumab and tremelimumab in humanized CTLA-4/hFcyR mice. A) Cartoon illustration of humanized CTLA-4 mice crossed to hFcyR mice to generate a hCTLA-4/hFcyR colony for the in vivo evaluation of fully human anti-CTLA-4 antibody variants. B) Flow cytometry evaluation of endogenous murine CTLA-4 cell surface expression in WT C57BL/6 mice compared to hCTLA-4/hFcyR mice, expressed as mean fluorescence intensity (MFI). C) Ipilimumab is a human IgG1 antibody whereas tremelimumab is a human IgG2 subclass. D) Surface plasmon resonance characterization of binding to human CTLA-4 by recombinant variants of ipilimumab and tremelimumab and their respective KDs. E) Evaluation of the ability of 10D1 and 1121 Fabs on IgG2 backbone to block interaction of hCTLA-4 with B7.1 as measured by ELISA. F) In vivo evaluation of the MC38 colon carcinoma tumor microenvironment of subcutaneous tumors following systemic (i.p.) administration of 10D1-IgG1 or 10D1-IgG2 when compared to IgG1 isotype control. Tumor bearing hCTLA-4/hFcγR mice were treated with 200 µg of anti-CTLA-4 mAb variants on days 0 and 3 following randomization, and sacrificed 24 hours later for flow cytometry analysis of CD8 T cells, CD4 effector T cells, and T regulatory cells (Tregs). G) Evaluation of the in vivo activity of anti-CTLA-4 subclass variant on MC38 tumor growth. Ipilimumab and tremelimumab were tested as both IgG1 and IgG2 Fc subclass variants. Tumor bearing hCTLA-4/hFcγR mice were treated with 200 µg of anti-CTLA-4 mAb variants on days 0, 3 and 6. H) Ipilimumab and tremelimumab were both placed on an IgG2 Fc backbone and tested for antitumor activity in the MC38 tumor model. I) In vivo evaluation of the B16 melanoma tumor microenvironment of subcutaneous tumors following systemic (i.p.) administration of 10D1-IgG1 or 10D1-IgG2 when compared to IgG1 isotype control. Tumor bearing hCTLA-4/hFcγR mice were treated with 200 µg of anti-CTLA-4 mAb variants on days 0 and 3 following randomization, and sacrificed 24 hours later for flow cytometry analysis of CD8 T cells, CD4 effector T cells, and T regulatory cells (Tregs). J) Ipilimumab in either and IgG1 or IgG2 format was tested for antitumor activity in the B16 tumor model. n = 4-8 mice per group; bars represent SEM, and *p < 0.05, **p < 0.01, ***p < 0.001. ****p < 0.0001.

### The inhibitory receptor FcyRIIB is upregulated in the tumor microenvironment (TME)

Given limitations of Treg depletion from anti-CTLA-4 antibodies in hCTLA-4/hFcyR mice, and lack of persistent Treg depletion in patients,^7^ we went back to the humanized model to identify potential mechanisms of resistance to antibody-mediated depletion in solid tumors. The ability of an effector cell to deplete a target coated with antibody depends on the collective balance of signaling through activating and inhibitory FcyRs, resulting in an activating/inhibitory (A/I) ratio. In mice, the IgG2a subclass is the most efficient ADCC effector because of its A/I ratio of 69:1.^9^ In humans, the ability of IgG1 or IgG2 subclasses to engage activating receptors is not nearly as strong (**Figure 2A**). We assessed the relative contribution of each activating and inhibitory FcyR in the TME of hCTLA-4/hFcyR mice. Here, we found that indeed the sole inhibitory receptor FcyRIIB is significantly upregulated on myeloid cells compared to the individual activating FcyRs (**Figure 2B**). This is important as it suggests that excessive FcyR inhibitory tone in the TME could act as another “checkpoint” blocking Treg depletion. Thus, one possible hypothesis is that the ratio of activating to inhibitory FcyRs in the tumor microenvironment further limits ADCC and could account for the lack of Treg depletion in humans treated with anti-CTLA-4 antibodies. We therefore characterized the expression of FcyRs in human tumors using quantitative multiplex immunofluorescence (mIF) to profile the expression of FcyRIII (CD16) and FcyRIIB (CD32b) on intratumoral CD68^+^ myeloid cells in oral squamous cell carcinoma biopsy specimens (**Figure 2C, Supplementary Table 1**), finding a mean CD68^+^ myeloid cell FcyRIIB expression of greather than 60%, resulting in a notably depressed intratumoral A/I ratio (mean < 2:1; range 0.5 to 4.0) (**Figure 2D)**.

**Figure 2.**
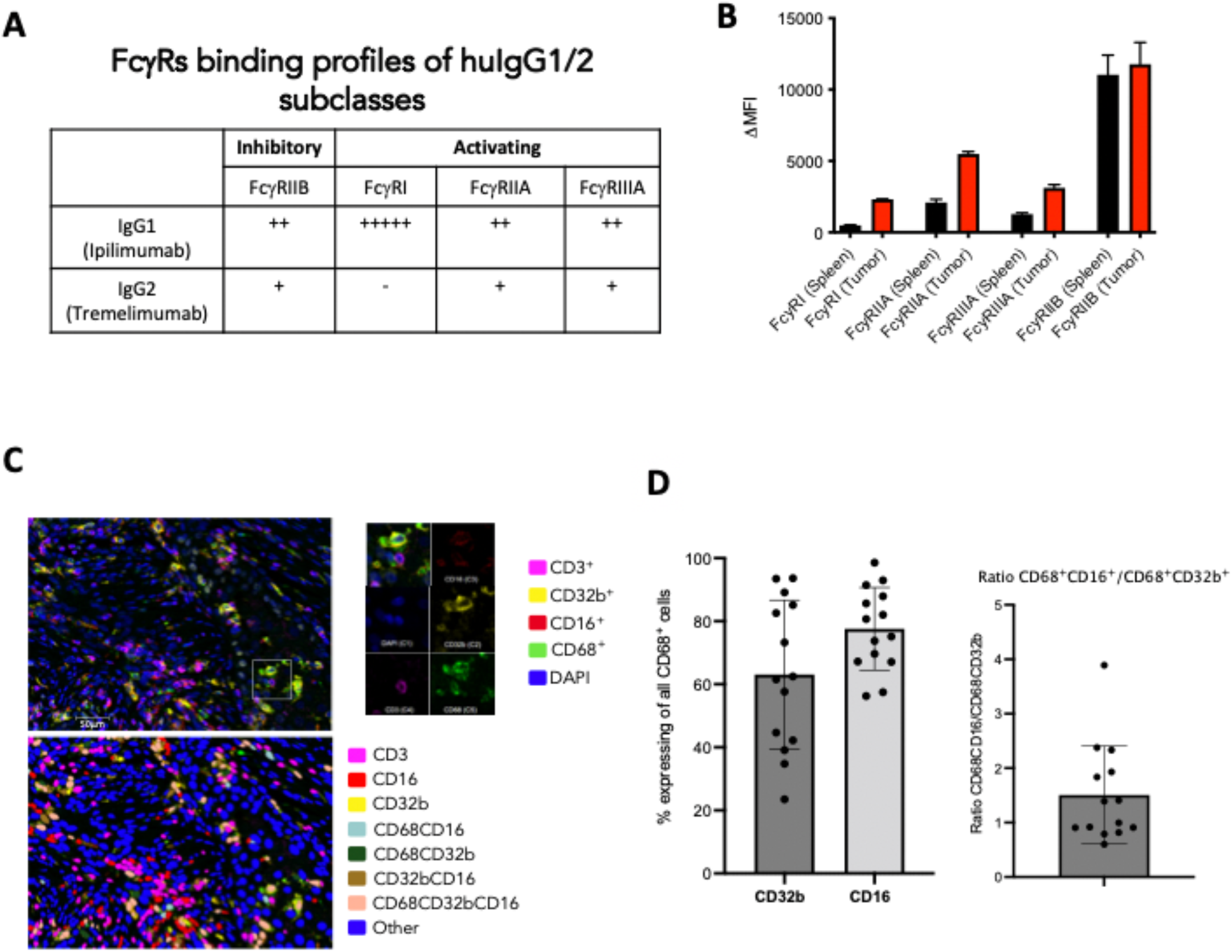
Murine and human tumor express high levels of FcyRIIB in the tumor microenvironment. A) Table demonstrating the relative binding affinities of ipilimumab or tremelimumab subclasses for activating and inhibitory FcyRs. B) Characterization of human FcyR expression in spleen or tumor infiltrating leukocytes. Each hFcyR was evaluated using flow cytometry of dissociated tissues. C) mIF of human tissue samples for hFcyRIIIA (CD16) and FcyRIIB (CD32B) on infiltrating myeloid cells using the PhenoImager HT. D) Quantification of CD32B and CD16 expression on tumor infiltrating CD68+ myeloid cells and ratio of staining of the activating (CD16) and inhibitory (CD32B) profile.

### Abrogating FcyRIIB engagement in the TME improves Treg depleting potency of anti-CTLA-4 antibodies and enhances anti-tumor immunity

Based on these data, we assessed if limiting binding to FcyRIIB could release this checkpoint and allow Treg depletion in our humanized mice. We first tested the ability of an FcyRIIB blocking antibody (clone 2B6) to improve the in vivo activity of anti-CTLA-4 antibodies. Here, co-administration of anti-FcyRIIB antibody in combination with the IgG1 version of Ipilimumab in the MC38 model led to significantly improved Treg depletion, expansion of CD8 and CD4 T cells(**Figure 3A**), as well as complete tumor control in all the treated mice (**Figure 3B**). Using ipilimumab (10D1) on an IgG2 Fc backbone, did not have a significant effect, as expected, given that IgG2 antibodies do not engage activating FcyRs at baseline and are poor at mediating Treg depletion through ADCC.

**Figure 3:**
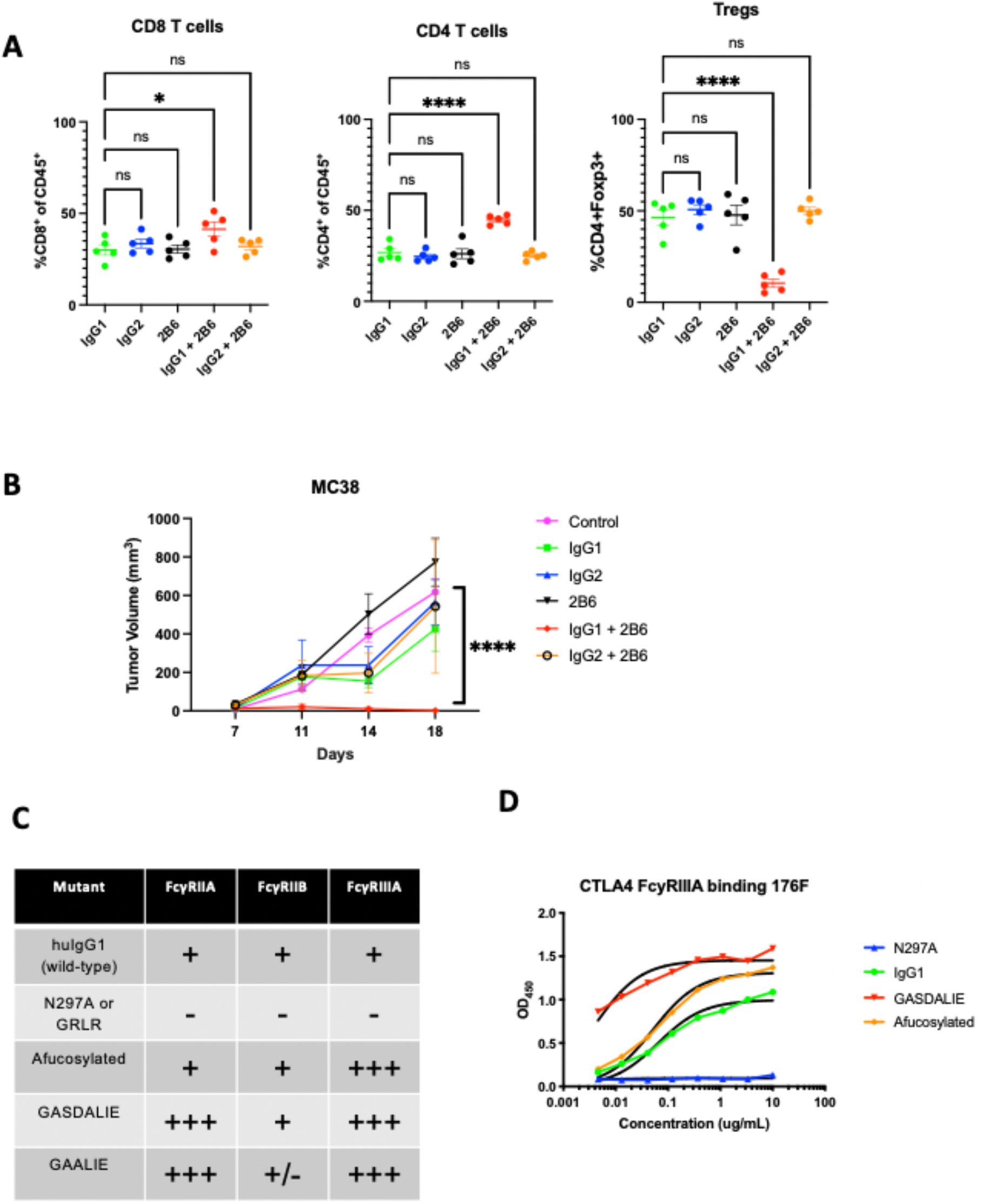
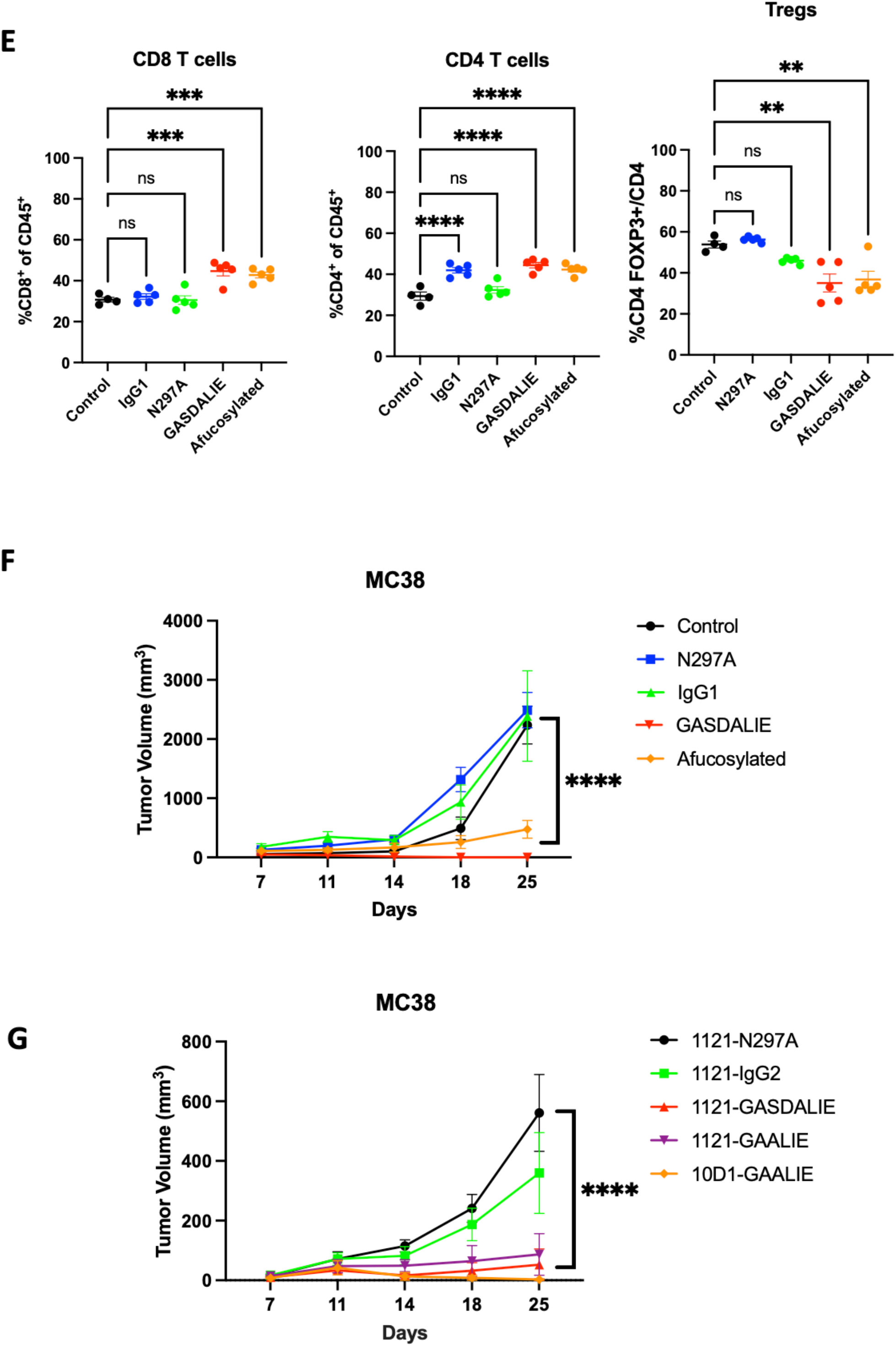
Limiting binding to FcyRIIB leads to enhanced Treg depletion and tumor control in vivo. A) In vivo evaluation of the effect of FcyRIIB blocking antibody (clone 2B6) on Treg depletion. Flow cytometric evaluation of the MC38 colon carcinoma tumor microenvironment following systemic (i.p.) administration of 10D1-IgG1 or 10D1-IgG2 (200 ug alone or in combination with 50 ug intratumoral 2B6 blockade prior on days 0 and 3) comapred to 2B6 only control. Mice were sacrificed 24 hours later and flow cytometry analysis was performed for CD8 T cells, CD4 effector T cells, and T regulatory cells (Tregs). Blockade of the inhibitory FcyR (FcyRIIB) leads to significantly less Tregs when coadministered with 10D1-IgG1 compared to 10D1-IgG2. B) In vivo evaluation of the effect of FcyRIIB blocking antibody (clone 2B6) on tumor control in the MC38 tumor model. Here, 200 ug of 10D1 subclass variants (e.g. IgG1 or IgG2) were given alone or in combination with 2B6 antibody blockade (50 ug given intratumoral) on days 0, 3, and 6. The combination of the IgG1-subclass variant led to significantly improved tumor control when compared to anti-CTLA-4 antibody alone or the anti-CTLA-4 IgG2 subclass antibody with 2B6. C) Table of the 10D1 IgG subclass variants generated for these studies, along with their relative binding affinities for activating vs inhibitory FcyRs. D) ELISA-based quantification of the relative affinity of the various subclass variants tested (in E and F). E) In vivo evaluation of the effect of Fc-enhanced anti-CTLA-4 antibodies (clone 10D1) on Treg depletion. Flow cytometric evaluation of the MC38 colon carcinoma tumor microenvironment following systemic (i.p.) administration of anti-CTLA-4 antibody variants (200 ug on days 0 and 3) comapred to to IgG control. Mice were sacrificed 24 hours later and flow cytometry analysis was performed for CD8 T cells, CD4 effector T cells, and T regulatory cells (Tregs). F) In vivo evaluation of the effect of Fc-enhanced anti-CTLA-4 antibodies (clone 10D1) on tumor control in the MC38 tumor model. Here, 200 ug of 10D1 subclass variants were given on days 0, 3, and 6. The Fc-enhanced variants led to significantly improved tumor control when compared to control, Fc-null (N297A), or IgG1. G) In vivo evaluation of the effect of Fc-enhanced anti-CTLA-4 antibodies (clone 10D1 or 1121) on tumor control in the MC38 tumor model. Here, 200 ug of 10D1 subclass variants were given on days 0, 3, and 6. The Fc-enhanced GAALIE variants led to significantly improved tumor control, even when placed on the 1121 (tremelimumab) backbone. n = 4-7 mice per group; bars represent SEM, and *p < 0.05, **p < 0.01, ***p < 0.001. ****p < 0.0001.

Monoclonal antibodies blocking the inhibitory FcyRIIB have been developed and tested in patients, but co-administration with anti-CTLA-4 antibodies is likely not feasible due to the large sink for FcyRIIB on circulating B cells and the potential toxicity of enhancing serum IgG triggered inflammation. Because binding of IgG antibodies to FcyRs is driven largely by their amino acid sequence (IgG1 vs IgG4) or composition of a conserved glycan at amino acid position 297^14^, we employed Fc-engineering to modify ipilimumab for an enhanced A/I ratio with the goal of enhancing its in vivo activity. The first variant, GASDALIE, is genetically modified to bind more strongly to activating FcyRs (FcyRIIA and FcyRIIIA). The second variant, afucosylated ipilimumab, is a “glyco-engineered” version in which removal of a core fucose on the conserved glycan enhances binding to FcyRIIIA approximately 10 to 20-fold, while retaining binding to FcyRIIB (**Figure 3C**). We demonstrated that each of these variants indeed increases affinity to FcyRIIIA (CD16) (**Figure 3D**), and this led to enhanced Treg depletion (**Figure 3E**) with significantly improved tumor control in hCTLA-4/hFcyR mice (**Figure 3F**). Swapping the Fc of tremelimumab (IgG2; 1121) with GASDALIE, or a recently described Fc variant from our group with an even more favorable A/I ratio and pharmacokinetic profile (-GAALIE),^18^ improved the in vivo activity of this anti-CTLA-4 antibody, while the ipilimumab 10D1-GAALIE construct showed the greatest activity (**Figure 3G**). These data define the inhibitory receptor FcyRIIB as a significant brake on Treg depleting antibodies in the TME, and highlight the importance of Fc-engineering for generating enhanced anti-CTLA-4 antibodies with improved in vivo activity.

### Potent targeting of Tregs with Fc-optimized anti-CCR8 antibodies

While CTLA-4 is upregulated on Tregs, it is also expressed on non-Treg subsets, both on the surface and in intracellular stores which can be mobilized following T cell activation. Blocking CTLA-4 on effector T cells is thought to be a primary driver of immune-related adverse events (irAEs), which occur in a substantial proportion of patients receiving anti-CTLA-4 antibody therapy. Similarly, antibodies to other targets such as CCR4 and CD25 have previously been tested for their ability to deplete Tregs in patients, showing similarly high levels of expression on non-Treg populations.^19^ In contrast, the G-protein coupled receptor CCR8 has been identified as highly expressed on tumor infiltrating Tregs, with minimal expression on T cells in peripheral tissues. ^19,20^ To test whether Fc-enhanced anti-CCR8 antibodies can lead to optimal Treg depletion and improved in vivo activity, we generated variants of a murine anti-CCR8 antibody (clone SA214G2) and grafted this onto Fc constructs with variable A/I ratios (GRLR [Fc null], IgG1, and GAALIE). We found that, in contrast to the unmodified IgG1 version, the anti-CCR8 GAALIE variant led to consistent Treg depletion across several syngeneic tumor models (**Figure 4A**), including the more aggressive MB49 bladder tumor model. We observed impressive and consistent single agent anti-CCR8 GAALIE enhanced tumor control in subcutaneous MC38, MB49 and B16 models (**Figure 4B**). Additionally, direct comparison of anti-CCR8 and anti-CTLA-4 targeting in the MC38 model, confirmed the superior activity of the Fc-enhanced anti-CCR8 GAALIE (**Figure 4C**). While both afucosylated anti-CTLA-4 and anti-CCR8 improved tumor control when compared to their parental IgG1 counterparts, each were further significantly enhanced by the -GAALIE mutation, characterized by decreased FcyRIIB binding.^18^

**Figure 4:**
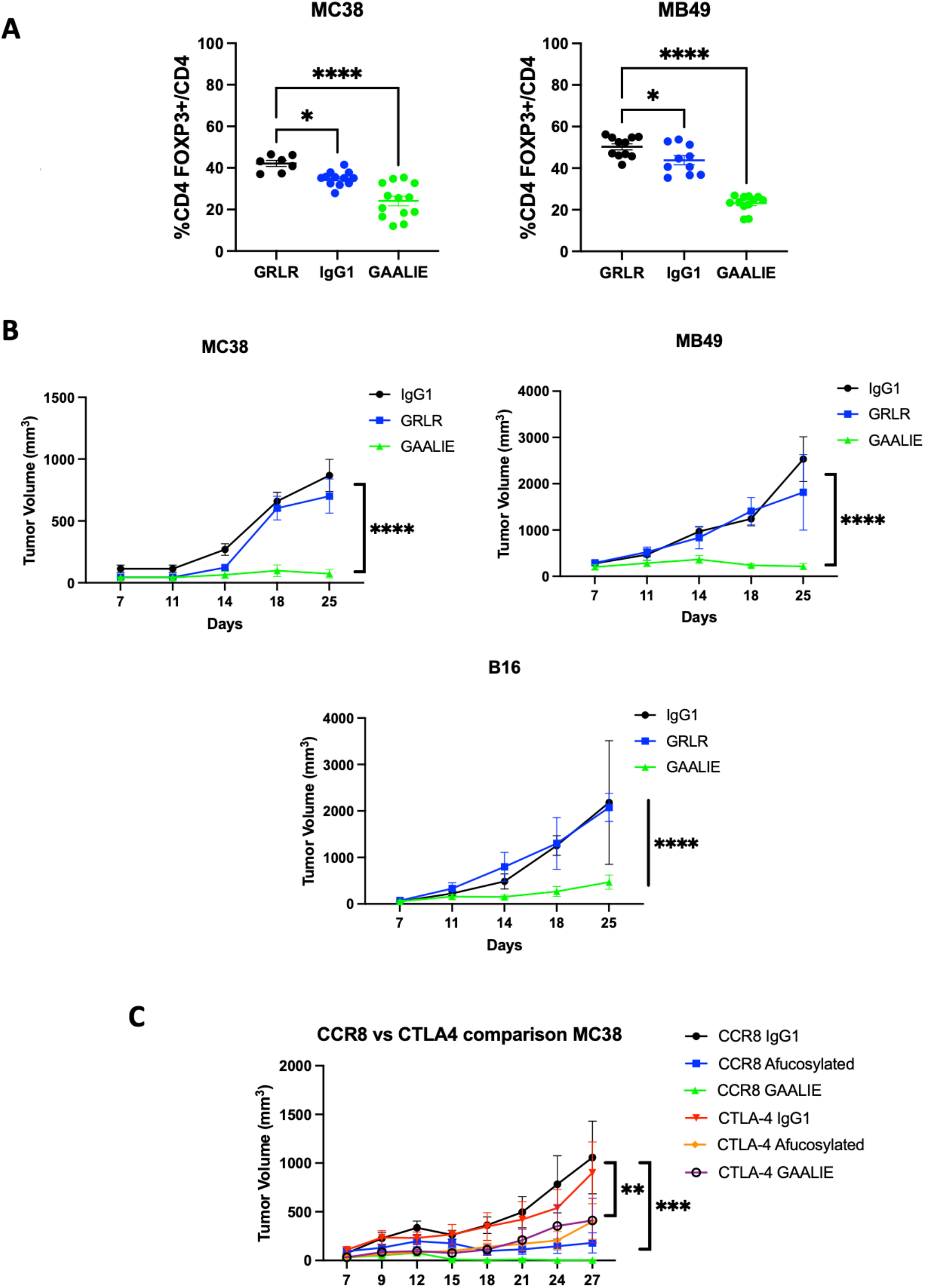
Potent targeting of Tregs with Fc-optimized anti-CCR8 antibodies. A) In vivo evaluation of the Fc-optimized anti-CCR8 antibody (GAALIE variant with decreased FcyRIIB binding) on Treg depletion. Flow cytometric evaluation of the MC38 colon carcinoma and MB49 bladder tumor microenvironment following systemic (i.p.) administration of 200 ug of CCR8-IgG1, CCR8-GRLR, or CCR8-GAALIE on days 0 and 3 following randomization. Mice were sacrificed 24 hours later and flow cytometry analysis was performed for CD8 T cells, CD4 effector T cells, and T regulatory cells (Tregs). B) In vivo evaluation of the effect of Fc-enhanced anti-CCR8 antibodies on tumor control in the MC38, MB49, and B16 tumor models. Here, 200 ug of anti-CCR8 subclass variants were given on days 0, 3, and 6. The Fc-enhanced variant GAALIE led to significantly improved tumor control when compared to Fc-null (GRLR), or IgG1. C) In vivo comparison of the effect of Fc-enhanced anti-CTLA-4 antibodies (clone 10D1) versus anti-CCR8 antibodies on tumor control in the MC38 tumor model. Here, 200 ug of each subclass variant was given on days 0, 3, and 6. The Fc-enhanced variants led to significantly improved tumor control when compared to IgG1 or afucosylated versions, with anti-CCR8 GAALIE demonstrating improved activity over anti-CTLA-4 GAALIE. n = 5-10 mice per group; bars represent SEM, and *p < 0.05, **p < 0.01, ***p < 0.001. ****p < 0.0001.

## Discussion

The approvals of antibodies blocking the inhibitory “checkpoints” PD-1 or CTLA-4 on T cells have offered new hope for durable remissions in patients with cancer. However, despite these encouraging advances our understanding of how these agents actually target and modulate immune subsets in humans remains incomplete. Prior preclinical studies have routinely demonstrated the potency of depleting T regulatory cells (Tregs) for optimal anti-tumor activity. For example, it has been previously demonstrated that in mice, the efficacy of anti-CTLA-4 antibodies was dependent on Fcy receptor (FcyR)-mediated Treg depletion, through engagement of myeloid cell activating FcyRIV to trigger antibody dependent cellular phagocytosis/cytotoxicity (ADCP/ADCC).^4^ In fact, without Treg depletion, the anti-tumor activity of anti-CTLA-4 antibodies in murine models is severely diminished.^21,22^ The first-generation human anti-CTLA-4 antibody ipilimumab is an IgG1 subclass antibody and would be predicted to drive ADCP/ADCC similar to other IgG1 antibodies like rituximab or trastuzumab. Initial studies in patients with melanoma supported the finding that Treg depletion in ipilimumab treated patients is associated with infiltrating CD16^+^ (FcyRIIIA) myeloid cells, with higher proportions of these cells in responding versus non-responding patients.^23^ Human FcyRIIIA, the homolog of murine FcyRIV, is one of the primary FcyRs contributing to ADCC/ADCP.^9^ Central to the mechanism of FcyR mediated depletion is the observation that engagement of activating FcyRs (hFcyRIIIA or mFcyRIV) are gated by co-engagement to the inhibitory FcyR, FcyRIIB,^9,10^ leading to a threshold for effector cell activation. When inhibitory receptor engagement is taken into account it is apparent that murine IgG2a, the most potent subclass for FcyR mediated cellular depletion, is over one log more potent that human IgG1, when cognate engagement of the relevant FcyRs is considered. Recent studies including patients across various solid tumor malignancies suggested that both ipilimumab and tremelimumab fail to decrease Tregs in human tumors,^7^ consistent with the decreased cytotoxic potential of the human IgG1 subclass as compared to its murine homologue. Attempts to overcome this species-specific limitation have focused on enhancing engagement to the activating Fc receptor FcyRIIIA (e.g. afucosylated antibodies) but have overlooked the potential role of FcyRIIB engagement as a limiting factor. The ability of these second generation FcyRIIIA enhanced anti-CTLA4 antibodies to deplete Tregs in human tumors has yet to be established.

To address the role of Fc receptors in mediating human anti-CTLA-4 antibody anti-tumor activity, we developed a fully humanized murine model allowing for the in vivo study of these antibodies in well established syngeneic tumor models (hFcyR/hCTLA-4 mice). Using this model, we found that the human anti-CTLA-4 antibodies (Ipilimumab, IgG1, Tremelimumab, IgG2) fail to routinely deplete Tregs in the TME and equally lack antitumor activity. Interestingly, this limitation appears to be due to an increase of the inhibitory FcyR (FcyRIIB) in the TME. Blocking FcyRIIB, or engineering anti-CTLA-4 antibodies with decreased binding to FcyRIIB, rescues Treg depletion and anti-tumor efficacy. These findings were further validated for antibodies targeting CCR8, a chemokine receptor with more selective expression than CTLA-4 on tumor-infiltrating Tregs.^19^ Collectively, these studies identify FcyRIIB as an important immune checkpoint in the solid tumor microenvironment and suggest Fc variant engineering as a direct means to overcome this limitation to anti-tumor immunity.

While our studies found that anti-CTLA-4-IgG1 antibodies can lead to Treg depletion in the immune responsive MC38 tumor model, this was not the case in the B16 melanoma tumor model. Additionally, regardless of Treg depletion neither IgG1 nor IgG2 anti-CTLA-4 antibodies lead to durable tumor control in either model. Although these data suggested that anti-CTLA-4 antibodies lead to transient Treg depletion, this was insufficient to drive durable anti-tumor immunity in either immune sensitive (MC38) or resistant (B16) tumor models, and failed to dramatically remodel the local immune microenvironment outside of Treg depletion.

Using an antibody to specifically block human FcyRIIB, ^13^ our studies demonstrated that this potently rescues in vivo Treg depletion in humanized mice and significantly enhances tumor control. Furthermore, we demonstrate that FcyRIIB is abundantly expressed by myeloid cells in the TME of human tumor specimens, resulting in a notably suppressed A/I ratio.

Preferable to co-administration of an FcRIIB blocking antibody, is the engineering of anti-CTLA-4 antibodies for enhanced engagement of activating FcyRs (FcyRIIA and FcyRIIIA), with reduced FcyRIIB binding (GAALIE) which leads to improved ADCC in several tumor models.^18^ In our current studies, anti-CTLA-4-GAALIE variants improved anti-tumor activity of both the ipilimumab and tremelimumab Fabs.

While CTLA-4 was one of the original immunotherapy targets and has revolutionized our approach to cancer therapy as a field, its expression is also found on a large number of peripheral and tissue-resident T cell populations, where its “immune checkpoint blockade” effects are felt to drive the high incidence of immune-related adverse events. Thus, many groups have worked to define alternative Treg targets, of which CCR8 has been shown to be more clearly differentially expressed on tumor infiltrating Tregs.^19,20^ Thus, to enhance the rigor of our approach to Fc-engineering Treg depleting antibodies by limiting FcyRIIB engagement, we tested anti-CCR8 antibodies in several tumor models. We found that Fc-enhanced anti-CCR8 antibodies generally outperformed anti-CTLA-4 antibodies in both their Treg-depleting capacity as well as tumor control, with the anti-CCR8-GAALIE variant leading to impressive antitumor immunity across multiple aggressive models including the B16 melanoma tumor model and the MB49 bladder tumor model.

Collectively, these studies directly apply to ongoing clinical trials testing afucosylated (BMS, NCT03110107) or amino acid sequence variants (DLE variant, Agenus, NCT03860272) of anti-CTLA-4 antibodies. These data would suggest that without further limiting engagement to the inhibitory FcyR, they will also not effectively lead to Treg depletion in patients.

## Methods

### Study Design

The primary objective of this study was to determine the anti-tumor activity and underlying mechanisms associated with antibodies targeting cell surface molecules on T regulatory cells. These studies involved controlled laboratory experiments using humanized subcutaneous implantable mouse models of cancer to assess the Fc-dependence of Treg depleting antibodies. Primary outcomes for most studies included tumor burden (measured by caliper) and survival. Tumor-bearing mice were either randomized into treatment groups, or baseline tumor burden was measured and groups were assigned with matching tumor burden between experimental conditions. Sample sizes were determined based on initial pilot experiments to estimate likely effect sizes and variation. For tumor growth studies, we expect that at least 5 mice/group will provide 80% power when setting alpha at 5%. Data were analyzed using the Grubb’s test for statistical outliers using an α value of 0.01. Data are representative of at least 2 experiments. Tumor measurements and survival monitoring were blinded.

### Mouse strains

All mice were bred and housed at Rockefeller University Comparative Bioscience Center under specific pathogen-free conditions. All experiments were performed in compliance with institutional guidelines and were approved by the Rockefeller University Institutional Animal Care and Use Committee (IACUC). C57BL/6J (WT) (Stock# 000664) mice were purchased from The Jackson Laboratory. Humanized mice that are fully backcrossed to the C57BL/6J background expressing human Fc receptors (FcgRa^null^, FcgRI^+^, FcgRIIa^R131+^, FcgRIIb^+^, FcgRIIa^F158+^, FcgRIIIb^+^) were generated and extensively characterized as previously described.^16^ hCTLA-4/hFcyR mice were generated by backcrossing a fully hCTLA-4 KI mouse^6^ to hFcyR mice. All mice used were 7-12 weeks old at time of experiment. All experiments used a mix of male and female mice.

### Cell Lines and Cell Culture

The murine tumor cell lines MC38, MB49, and B16 were cultured inside tissue culture treated flasks with DMEM with 10% FCS (Hyclone) and 1X penicillin-streptomycin (100 units/ml, Gibco) at 37C and 5% CO_2_. Cells were split twice per week and cell viability was measured using trypan blue staining in the Countess II automated cell counter (Thermo Fisher).

### Generation of anti-human CTLA-4 and anti-CCR8 Abs and their Fc variants

The variable heavy and light regions of ipilimumab (10D1) or tremelimumab (1121) were synthesized based on their published sequences. The variable heavy region of anti-mouse CCR8 (Clone SA214G2, Biolegend) was determined by Mass Spectrometry (Bioinformatics Solutions, Inc.). The anti-human FcyRIIB (clone 2B6) was previously generated in our lab (Rockefeller University) and produced by the MSKCC Hybridoma Facility. The variable region sequences of the parental Abs were cloned and inserted into mammalian expression vectors with human IgG1 or human kappa Fc backbones. For the generation of human IgG1 N297A, GRLR, GASDALIE, and GAALIE Fc-domain variants, site-directed mutagenesis using specific primers was performed (Agilent Technologies) according to the manufacturer’s instructions. Mutated plasmid sequences were validated by direct sequencing (Azenta Biosciences). To produce antibodies, antibody heavy and light chain expression vectors were transiently transfected into Expi293 suspension cells (ThermoFisher). For the generation of the human IgG1 afucosylated Fc-domain variant, 200µM of 2-Deoxy-2-fluoro-L-fucose (Biosynth Carbosynth) was added 1 day after transfection. The secreted antibodies in the supernatant were purified by protein G Sepharose 4 Fast Flow (GE Healthcare). Purified antibodies were dialyzed in PBS and sterile filtered (0.22 μm). Purity was assessed by SDS-PAGE and Coomassie staining. Percentage of afucosylated forms was assessed by mass spectrometry.

### ELISA

Binding specificity, affinity, and blocking activity of anti-CTLA-4 monoclonal antibodies were determined by ELISA using recombinant human B7.1 (Sino Biological, #10698-HCCH), human CTLA-4 (Sino Biological, 11159-HNAH), and human FcyRs (human FCGR2A #10374-H08H, human FCGR2B #10259-H08H, human FCGR3A #10389-H08H, Sino Biological). 96 well ELISA Half Area High Binding plates (Greiner Bio-One, #675061) were coated overnight at 4°C with recombinant CTLA-4 proteins (1µg/mL) or human FcyRs (2µg/mL). All sequential steps were performed at room temperature. After washing, the plates were blocked for 1 hour with 1xPBS containing 2% BSA (for CTLA-4 or B7.1/B7.2 proteins) or 10% BSA (for human FcgRs), and were subsequently incubated for 2 hours with serially diluted IgGs (dilutions are indicated in the figures and were prepared in the relevant blocking solution). After washing, plates were incubated for 1 hour with horseradish peroxidase conjugated anti-human IgG (#109-035-088, Jackson IummunoResearch). For ELISA inhibition assay, after blocking non-specific sites, plates were incubated for 1 hour with serially diluted IgGs with 1µg/mL of human CTLA-4-Biotin (Sino Biological, #11159-H08H-B) in 1xPBS with 2% BSA. After washing, plates were incubated for 1 hour with horseradish peroxidase labeled streptavidin (#405210, BioLegened). Detection was performed using a one component substrate solution (TMB), and reactions stopped with the addition of 0.18 M sulphuric acid. Absorbance at 450 nm was immediately recorded using a SpectraMax Plus spectrophotometer (Molecular Devices), and background absorbance from negative control samples was subtracted.

### Subcutaneous Tumor Implantation

MC38, MB49, and B16 cells were detached using trypsin digestion from tissue culture flasks and assessed for viability (>95%). Cells were washed thrice with PBS and were resuspended in sterile cold PBS. Mice were injected in their lower abdominal flanks with 100 ul of this mixture, corresponding to 2 × 10^5^ (for MB49 and B16) and 2 × 10^6^ (for MC38) cells inoculated per mouse flank. Tumors were measured 2-3 times per week using an electronic caliper. Volume was calculated using the formula (L_1_^2^ x L_2_)/2, where L_1_ is the longest dimension. Tumors were randomized into treatment groups when tumors were approximately 5-6 mm in diameter. Mice were randomized by tumor size (day 0), and received treatment as described in the legend of each experiment. hCTLA-4/hFcγR mice were treated with 200 µg of anti-CTLA-4 mAb variants, 50 µg of anti-human FcyRIIB or control PBS (intratumoral) on days 0, 3 and 6. Mice were monitored after treatment initiation and sacrificed when tumor size reached the Rockefeller University IACUC limit.

### Flow Cytometry

Tumor tissues were digested using Mouse Tumor Dissociation kits (Miltenyi) according to the manufacturer’s protocols. Isolated cells were stained for viability using the Aqua Amine fixable live dead dye (Thermo Fisher) at room temperature using standard protocols. After viability staining, cells were resuspended in FACS buffer (PBS with 0.5% BSA and 2 mM EDTA) and Fc blocked using human TrueStain FcX (Biolegend). For cell surface staining, cells were incubated for 30 minutes at 4C with the following anti-mouse antibodies from Biolegend CD4-FITC, clone GK1.5, CD8a-BV786, clone 53-6.7; CD3e-Percp-Cy5.5, clone 17A2; and ThermoFisher CD45-AlexaFluor 700, clone 30-F11. Cells were then washed twice, fixed, and permeabilized (eBiosciences Intracellular Fixation & Permeabilization Buffer Set) prior to intracellular staining with FOXP3-PE, clone 150D. Samples were analyzed using an Attune NxT flow cytometer (Thermo Fisher) and data was analyzed using FlowJo 10 (TreeStar).

### Human tumor tissues

Baseline oral squamous cell carcinoma 4mm punch biopsy samples were obtained between November 2016 and August 2019 from consenting patients planned for standard of care surgical resection, under a Providence Health & Services IRB approved clinical study protocol (IRB #16-042) at the Earle A. Chiles Research Institute, Providence Portland Medical Center, Oregon USA.

### Ethics statements

The study was approved by the Providence Health & Services Institutional Review Board (IRB# 16-042). Patient consent was obtained as a pre-requisite for study enrollment. Patient consent for publication is not required.

### Multiplex Immunofluorescence staining (mIF)

To prepare specimens for multiplex IF staining, tissue sections were cut at 4 µm from formalin-fixed paraffin-embedded blocks. All sections on slides were deparaffinized using the Bond ER2 Leica Biosystems AR9640, followed by staining with the Leica Bond RX autostainer. The slides were stained with - CD3 (1:50 SP7 Abcam ab16669), CD16 (1:50 2H7 CellSignaling Technologies 88251S), CD32b (1:2000 NovusBiologicals NBP2-89364), and CD68 (1:400 PGM-1 DAKO). Antibody binding was visualized using secondary anti-mouse/anti-rabbit polymer HRP followed with TSA-Opal reagents (Akoya NEL811001KT). Tissue slides were incubated with DAPI as a counterstain and coverslipped with VectaShield mounting media (Vector Labs). Control tissue samples were stained for each marker as positive controls.

Whole slide digital images were captured with the PhenoImager HT platform and analyzed with QuPath software^24^. Tissue samples from consecutive slides were stained by conventional H&E and scanned with the Leica SCN400F platform. H&E slides in conjunction with Cytokeratin staining were used to annotate tumor regions within the whole slide image. Cells were segmented, phenotyped, and enumerated using the QuPath object classification algorithm.

### Data and statistical analysis

For murine studies, data was analyzed using Prism GraphPad software. One-way ANOVA with Tukey’s multiple comparison test was used to compare 3 or more groups. Comparisons between 2 groups used unpaired two-tailed t tests for statistical significance. All data, unless otherwise indicated, are plotted as mean ± standard deviation (s.d.). Kaplan-Meier curves used Log-Rank tests to calculate statistical significance. For all statistical tests, * p < 0.05, ** p < 0.01, *** p < 0.001, **** p < 0.0001, a p value under 0.05 was considered statistically significant, not significant values are denoted as n.s. Lines associated with asterisks indicate groups compared for significance.

## Acknowledgements

The authors would like to members of the J.V.R. laboratory for excellent technical assistance and helpful feedback. This work was supported by following grants from the National Institutes of Health (NIH): UL1TR001866 and KL2TR001865 from the National Center for Advancing Translational Sciences NIH Clinical and Translational Science Award Program, D.A.K.; K08CA248966 to D.A.K.; R01CA244327, R35CA196620, and P01CA190174 to J.V.R.; MSK Specialized Program of Research Excellence in Bladder Cancer P50CA221745 through a Developmental Research Program Award; and the National Cancer Institute Cancer Center Support Grant P30CA008748. The content is solely the responsibility of the authors and does not necessarily represent the official views of the NIH. C.B.M., S.J., R.L., and B.F. would like to thank the Harder Family, Robert W. Franz, Elsie Franz Finley, the Murdock Trust, the Chiles Foundation, Nancy Lematta. R.L. is also supported by Bristol Myers Squibb IO-ON.

## Author contributions

**Conceptualization**, D.K., R.L, B.F., and J.R.; **Methodology**, D.K., R.L., R.D.; **Validation**, D.K.,R.D., S.J., R.M., A.J. C.B-M., M.B., D.S., B.P., C.B.; **Formal Analysis**, D.K.,R.D., S.J., R.M., A.J. C.B-M., M.B..; **Writing**, D.K., R.L, B.F., and J.R.; **Resources**, D.K., R.L, B.F., and J.R.; **Supervision**, D.K., R.L, B.F., and J.R.

**COI:** The authors declare no financial conflicts of interests relevant to this work.

---

**Supplementary Figure 1:**
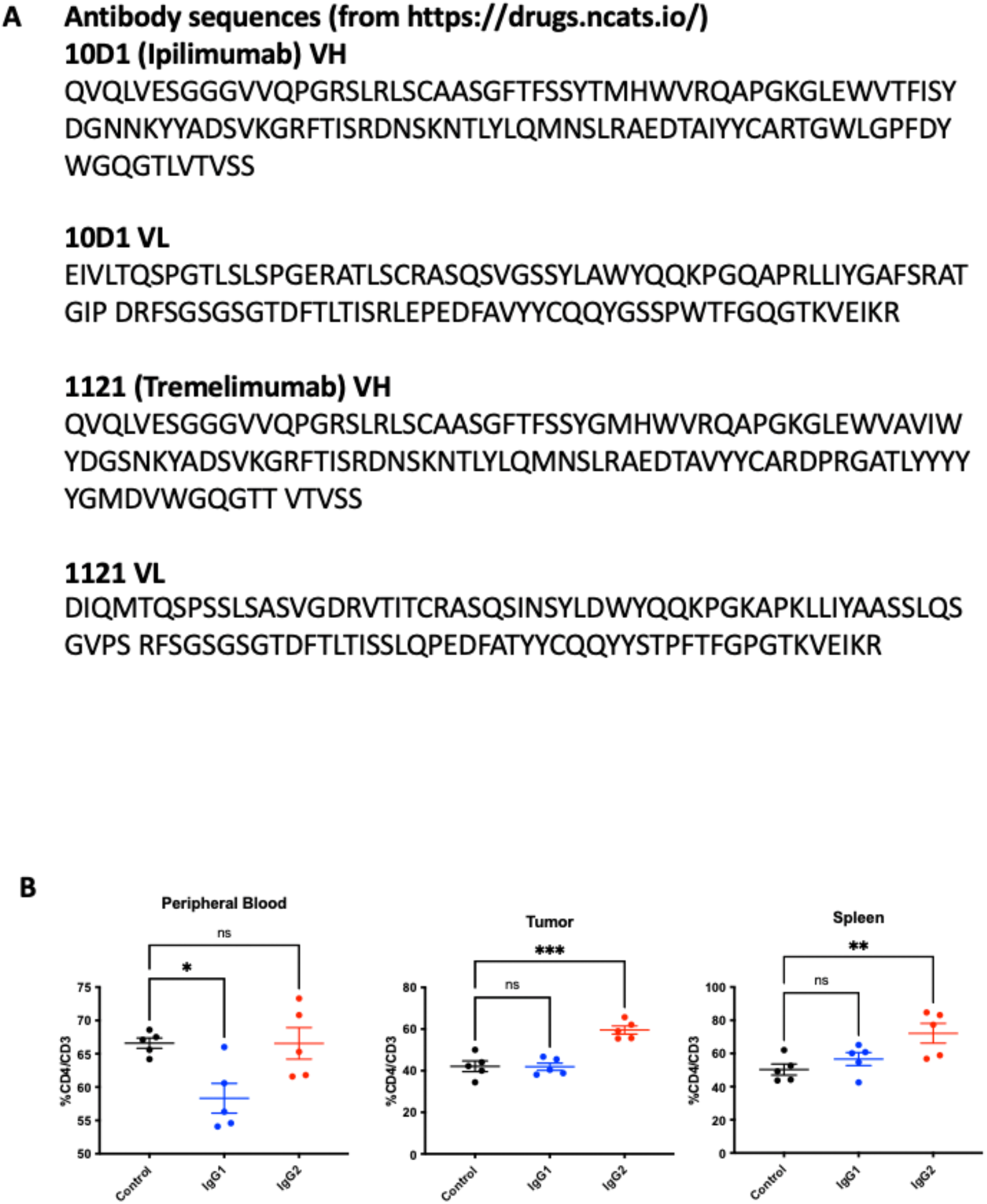
A) Amino acid sequence of each of the human anti-CTLA-4 antibodies used in these studies. B) In vivo evaluation of the MC38 colon carcinoma tumor microenvironment of subcutaneous tumors following systemic (i.p.) administration of anti-CD4-IgG1 or anti-CD4-IgG2 (both clone GK1.5) when compared to IgG1 isotype control. Tumor bearing hCTLA-4/hFcγR mice were treated with 200 µg of anti-CD4 mAb variants on days 0 and 3 following randomization, and sacrificed 24 hours later for flow cytometry analysis of CD4 T cells in the peripheral blood, tumor, or spleen.

**Supplementary Table 1.** Demographics of HNSCC patient tumor biopsies.

| Age | Gender | Stage | Smoker |
| --- | --- | --- | --- |
| 58 | M | T3 N3b | Yes |
| 68 | M | T4b N0 | Yes |
| 63 | F | T2 N0 | No |
| 52 | M | T3 N0 | Yes |
| 72 | F | T2 N2c | No |
| 66 | M | T3 N1 | Yes |
| 51 | M | T1 N0 | Yes |
| 52 | F | T1 N0 | No |
| 60 | M | T3 N2b | Yes |
| 66 | F | T2 N0 | Yes |
| 42 | F | T1 N0 | Yes |
| 41 | M | T1 N0 | No |
| 50 | M | T2 N0 | No |
| 64 | M | T1 N0 | No |
| 83 | M | T4a N0 | No |
| 74 | M | T3 N2a | Yes |
| 61 | M | T2 N1 | Yes |

